# The bHLH-PAS transcriptional complex Sim:Tgo plays active roles in late oogenesis to promote follicle maturation and ovulation

**DOI:** 10.1101/2022.12.30.522327

**Authors:** Rebecca Oramas, Elizabeth Knapp, Baosheng Zeng, Jianjun Sun

## Abstract

Across species, ovulation is a process induced by a myriad of signaling cascades that ultimately results in activation of proteolytic enzymes and degradation of the follicle wall to release encapsulated oocytes. Follicles need to first mature and gain ovulatory competency before ovulation. However, the signaling pathways regulating follicle maturation are incompletely understood in *Drosophila* and other species. Our previous work showed that bHLH-PAS transcription factor Single-minded (Sim) likely plays important roles for follicle maturation downstream of the NR5A-family nuclear receptor Ftz-f1 in *Drosophila*. Here, we explore the mechanism of Sim-regulated follicle maturation. We demonstrate that Tango (Tgo), another bHLH-PAS protein acts as a cofactor of Sim to promote follicle cell differentiation from stages 10 to 12. In addition, we discovered that re-upregulation of Sim in stage-14 follicle cells is also essential to promote ovulatory competency by upregulating octopamine receptor in mushroom body (OAMB), matrix metalloproteinase 2 (Mmp2), and NADPH oxidase (NOX), either independent of or in conjunction with the zinc-finger protein Hindsight (Hnt). All of these factors are critical for successful ovulation. Together, our work indicates that the transcriptional complex Sim:Tgo plays multiple roles in late-stage follicle cells to promote follicle maturation and ovulation.

## Introduction

Reproductive health is a vital component of every society. In the United States alone, 10% of reproductively mature women suffer from infertility (Chandra et al., 2014). Some of the leading causes of infertility are ovulation disorders, which affect 1 in 4 couples who have difficulty conceiving (Health (UK), 2013). Ovulation is the process of releasing a fertilizable oocyte from a mature ovarian follicle. In mammals, this process is induced by a luteinizing hormone (LH) surge which activates multiple pathways including progesterone, EGFR-Ras-ERK1/2, and prostaglandin signaling (Richards and Ascoli, 2018; Robker et al., 2018; Duffy et al., 2019; Tokmakov et al., 2020). Despite many years of investigation, we are still lacking detailed mechanisms regarding the precise regulation of follicle rupture, the actual step of releasing the oocyte.

*Drosophila melanogaster* offers a wide array of genetic tools and displays significant conservation of ovulation pathways with mammals, making it a valuable model to decipher the genetic pathways for follicle rupture. In *Drosophila*, the adrenergic system is responsible for inducing multiple pathways for follicle rupture and ovulation (Monastirioti et al., 1996; Cole et al., 2005; Middleton et al., 2006; Deady and Sun, 2015). Octopamine (OA), the Drosophila counterpart of norepinephrine, binds to its receptor octopamine receptor in mushroom body (OAMB) in mature follicle cells and increases intracellular Ca^2+^ (Deady and Sun, 2015). In posterior follicle cells, the increase of intracellular Ca^2+^ induces the activation of Matrix metalloproteinase 2 (Mmp2), which breaks down the posterior follicle wall to allow oocyte release (Deady et al., 2015). In mainbody follicle cells, the Ca^2+^ activates NADPH oxidase (NOX), which increases ROS production along with superoxide dismutase 3 (SOD3) to facilitate follicle rupture (Li et al., 2018). Although we are gaining a better understanding of the rupture process, it is largely unknown how these ovulatory factors are precisely regulated. In other words, how do follicles become fully mature and competent to ovulate?

The *Drosophila* ovary is composed of ∼16 ovarioles, where follicles develop through 14 stages from anterior to posterior (Spradling, 1993). To reach maturity, somatic follicle cells surrounding the germline cells must undergo 8-9 rounds of mitosis from stages 1-6, three rounds of endocycle from stages 7-10A, and synchronized gene amplification and chorion synthesis from stages 10B-14 (Klusza and Deng, 2011). Our previous work has shown that ecdysteroid signaling induces the transition from the endocycle to synchronized chorion gene amplification by upregulating Ttk69, while it induces the expression of NR5A nuclear receptor Fushi tarazu-factor 1 (Ftz-f1) at stage 10B to regulate follicle cell differentiation, maturation, and competency to ovulatory stimuli (Sun et al., 2008; Knapp et al., 2020). Specifically, Ftz-f1 directly activates another transcription factor Single-minded (Sim) at stages 10B-12, which mediates final differentiation of follicle cells, manifested by the downregulation of Ttk69, Cut, Broad-Complex (Br-C), Ecdysone receptor isoform A (EcR.A) and EcR.B1, and upregulation of zinc-finger transcription factor Hindsight (Hnt) (Knapp et al., 2019; Knapp et al., 2020). Hnt subsequently activates OAMB expression in all follicle cells and Mmp2 expression in posterior follicle cells at stage 14 (Deady et al., 2017). Previous work noted that Sim is downregulated at stage 13 and re-upregulated at stage 14 after its expression from stage 10B to 12 (Knapp et al., 2020). However, it is unknown why Sim has such a dynamic pattern in later stages and how Sim precisely regulates follicle cell differentiation and final maturation.

Sim is well studied during embryonic development, where it functions as a master regulator of CNS midline cell transcription and development (Nambu et al., 1990). It belongs to the basic-helix-loop-helix-PAS (bHLH-PAS) protein family and forms a transcriptional activator with another bHLH-PAS protein Tango (Tgo; Nambu et al., 1991; Ohshiro and Saigo, 1997; Sonnenfeld et al., 1997; Estes et al., 2001; Jiang and Crews, 2003). Sim’s role is not only limited to CNS midline development but also critical for left-right asymmetry of embryonic gut, axon guidance in larval brain, lamina-retina neuron association during optic lobe development, and genital disc development (Pielage et al., 2002; Umetsu et al., 2006; Maeda et al., 2007; Freer et al., 2011). Tgo can not only heterodimerize with Sim but also with other bHLH-PAS proteins including Trachealess and Spineless (Ohshiro and Saigo, 1997; Emmons et al., 1999). The process of dimerization is critical for Tgo nuclear localization (Ward et al., 1998), while the specific PAS domain confers the transcriptional specificity (Zelzer et al., 1997). Despite these studies, little is known about Sim and Tgo’s role in adult physiology and reproduction.

In this paper, we demonstrate that Tgo is co-expressed with Sim throughout late oogenesis and that its nuclear translocation depends on Sim, which is consistent with the previous report. Moreover, disruption of Tgo and Sim showed similar defects in follicle cell differentiation, dorsal appendage formation, and egg morphology, suggesting that Sim and Tgo form a heterodimer complex to regulate follicle cell differentiation. Most importantly, we characterize a novel role of Sim in stage-14 follicles that is distinct from its role in follicle cell differentiation from stages 10B to 12. In stage-14, Sim promotes ovulatory competency by upregulating the expression of key proteins required for follicle rupture including OAMB, NOX and Mmp2. In addition, Sim’s role in promoting ovulatory competency is either independent of or in conjunction with Hnt. Our results suggest that Sim acts as a master regulator to promote ovulatory competency of the follicles. We also demonstrated for the first time that the tunable CRISPR-Cas9 system (Port et al., 2020) can effectively disrupt gene expression in a tissue-specific manner in follicle cells with 16 copies of the genome. Altogether, our work demonstrated that Sim:Tgo complex plays essential roles to promote follicle differentiation and maturation in late oogenesis. Considering the conserved nature of this family of transcription factors in neurogenesis (Crews and Fan, 1999), the role of this complex in follicle development may be conserved in other species.

## Results

### Tgo is dynamically co-expressed with Sim in follicle cells during late oogenesis

Sim and Tgo form a heterodimer complex in regulating CNS midline cell development (Sonnenfeld et al., 1997). To test whether Tgo is also a heterodimerization partner for Sim in late oogenesis, we first examined the Tgo expression in *Drosophila* ovaries using an anti-Tgo antibody. Tgo is faintly detected in the cytoplasm of follicle cells in mid-stage egg chambers and upregulated at late stage 10A (Fig 1A and S1A-B). Tgo is then enriched in follicle cell nuclei from stages 10B to 12 but becomes downregulated at stage 13 (Fig 1B-D). At stage 14, Tgo is re-upregulated and enriched in follicle cell nuclei (Fig 1E). The specificity of the Tgo staining is confirmed by the observation that Tgo expression in follicle cells is significantly reduced upon *tgo* knockdown via RNA interference (RNAi; Fig S1A-F).

**Figure 1.**
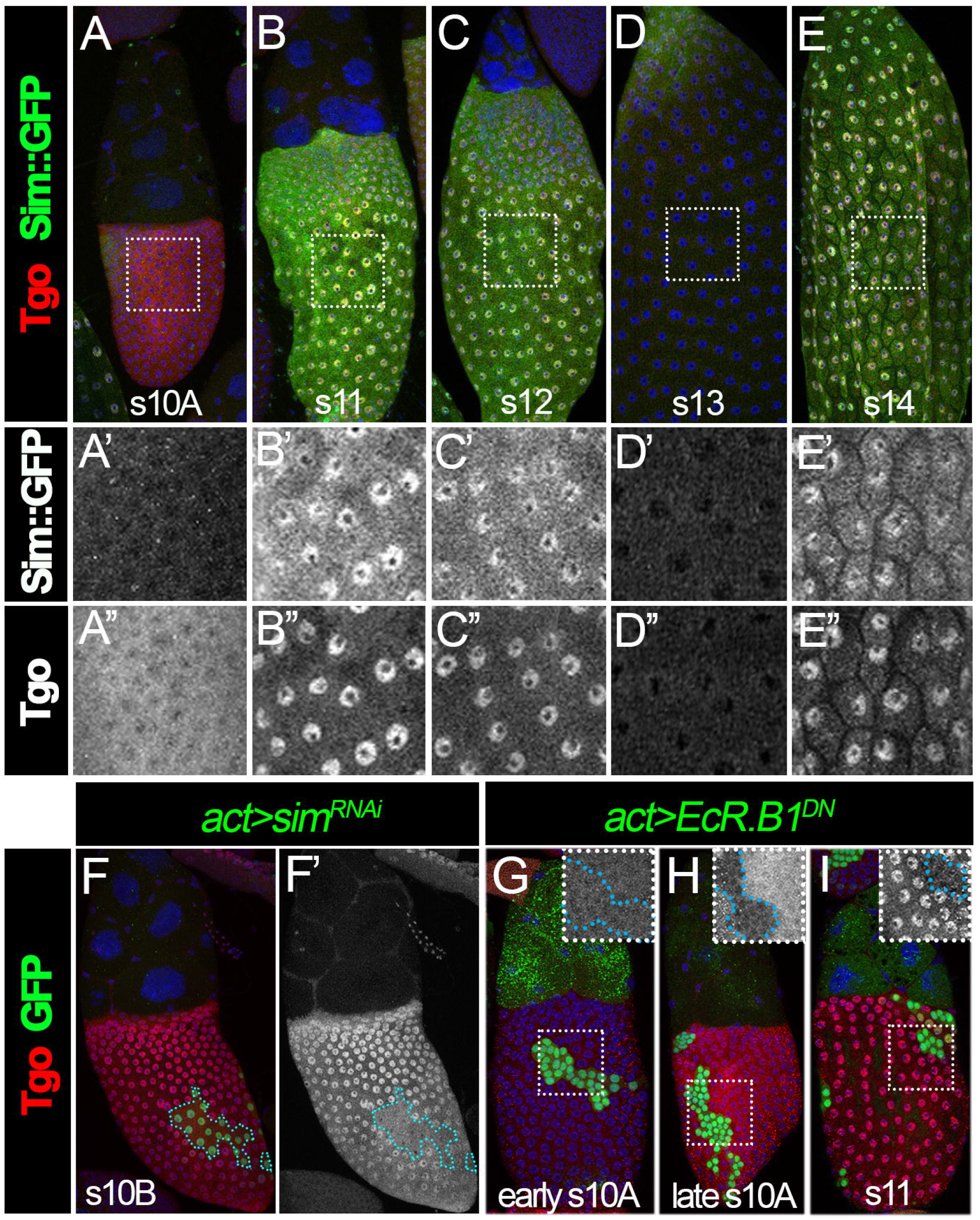
Tgo is dynamically expressed in follicle cells during late oogenesis. **(A–E)** Expression of Sim::GFP and Tgo in stages 10-14. Sim::GFP expression is detected by anti-GFP antibody (green) and Tgo is detected by anti-Tgo antibody (red) in stage-10A (A), stage-11 (B), stage-12 (C), stage-13 (D), and stage-14 (E) follicles. The higher magnification of the squared areas are shown in A’-E’ (Sim::GFP) and A”-E” (Tgo). All images from A-E were acquired using the same microscopic settings. **(F)** Tgo expression (red) in stage-10B follicles with flip-out Gal4 clones (green) overexpressing *sim*^*RNAi*^ (*act >sim*^*RNAi*^). (F’) Tgo expression (white) in wildtype vs. clone populations (outlined in blue). **(G-I)** Tgo expression (red) in stage-10A (G), stage-10B (H) and stage-11(I) follicles with flip-out Gal4 clones (green) overexpressing *EcR*.*B1*^*DN*^ (*act >EcR*.*B1*^*DN*^). Insets show higher magnification of Tgo expression in squared area. The clone boundary is outlined by a pink dashed line. Nuclei are marked by DAPI in blue.

The nuclear localization of Tgo coincides with the expression of Sim at stages 10B-12 and stage 14 (Fig 1B-E), which is consistent with our previous report (Knapp et al., 2020). Previous work showed that Tgo nuclear localization depends on Sim in CNS midline precursor cells (Ward et al., 1998). To test whether Sim is also required for Tgo nuclear localization in follicle cells, we examined the Tgo expression in *sim*-knockdown follicle cells. Tgo did not show nuclear enrichment but was rather maintained in the cytoplasm of *sim*-knockdown follicle cells (Fig 1F). This result confirms that Tgo nuclear localization depends on Sim in follicle cells and suggests that Tgo is likely the heterodimerization partner of Sim for its functions in late oogenesis.

We noticed an increase in Tgo expression at late stage 10A in comparison to earlier stages, which coincides with the endocycle-to-gene-amplification transition initiated by ecdysone signaling (Fig 1A; Sun et al., 2008). To test whether ecdysone signaling is required for the upregulation of Tgo expression, we examined the Tgo expression in follicle cells with overexpression of a dominant negative (DN) form of *EcR* (*EcR*^*DN*^). In earlier stages, *EcR*^*DN*^-overexpressing follicle cells showed the same level of Tgo expression as adjacent wild-type follicle cells (Fig 1G). However, we did not observe Tgo upregulation in *EcR*^*DN*^-overexpression follicle cells at late stage 10A, when the adjacent wild-type follicle cells had a significant increase of Tgo expression (Fig 1H). At stage 11, overexpression of *EcR*^*DN*^ prevented induction of Sim, which further blocks Tgo translocation into the nucleus (Fig 1I). This suggests that ecdysone signaling is required for the upregulation of Tgo expression at the stage 10A/10B transition.

### Tgo acts as a heterodimer partner of Sim for follicle cell differentiation

Disrupting the expression of Sim (as well as its upstream regulator Ftz-f1) prevents follicle cells from differentiating into a mature state at stage 14, which is characterized by the upregulation of Hnt and the downregulation of Cut, Ttk69, Br-C, EcR.A, and EcR.B1 (Knapp et al., 2019). To determine whether Tgo is indeed the partner for Sim in regulating follicle cell differentiation, we tested whether Tgo depletion affects the expression of some of these factors (Br-C, Cut, and Hnt) in stage-14 follicle cells. Consistent with previous findings, *sim*-knockdown follicle cells had extended expression of Br-C and Cut and reduced expression of Hnt at stage 14 (Fig 2A-C; Knapp et al., 2020). Similarly, *tgo*-knockdown follicle cells failed to downregulate Br-C and Cut (Fig 2D-E) and upregulate Hnt (Fig 2F) at stage 14. These results indicate that Tgo is also required to promote follicle cell differentiation and is likely the partner of Sim.

**Figure 2.**
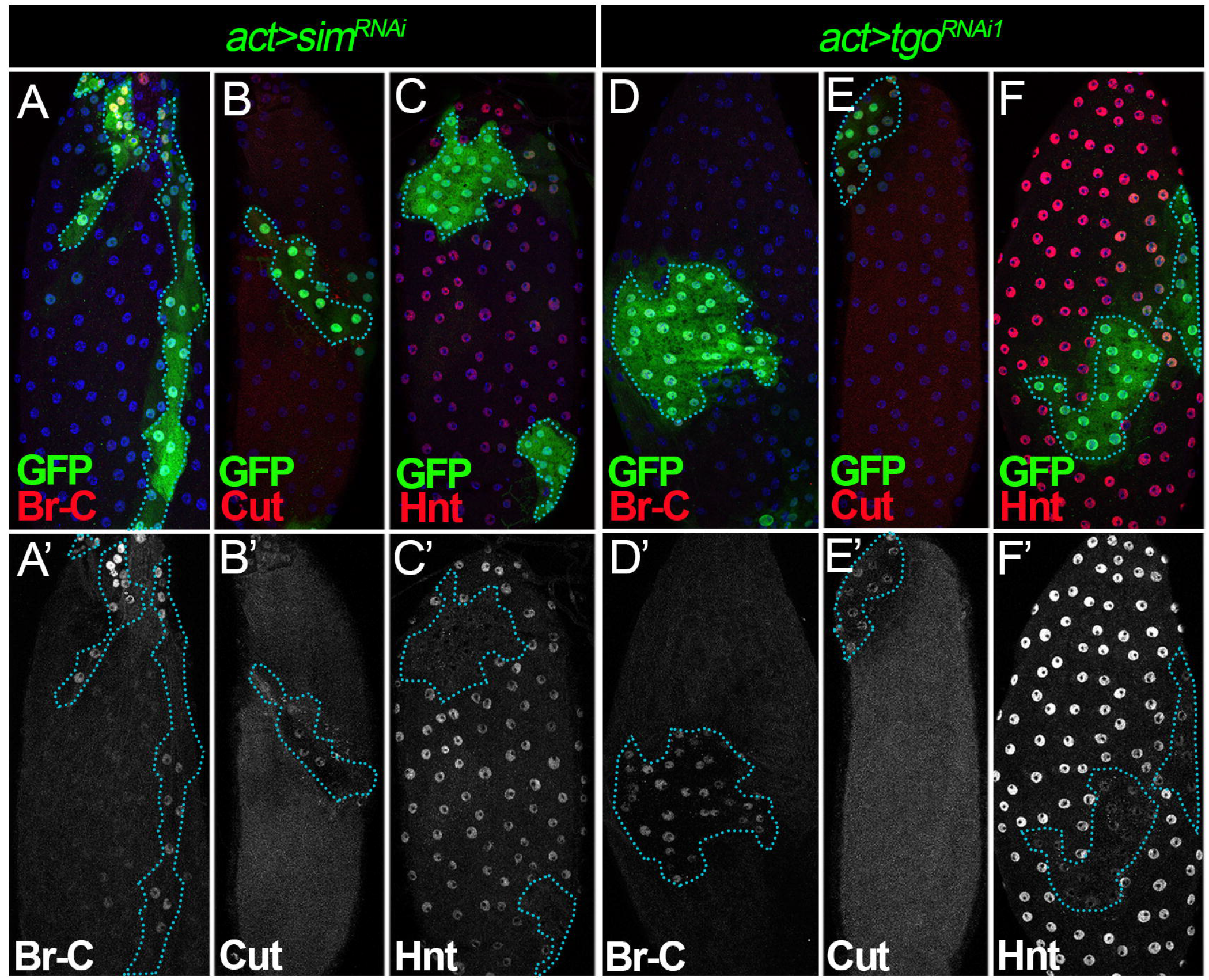
Tgo is a heterodimer partner of Sim for follicle cell differentiation. **(A–C)** Br-C (A), Cut (B) and Hnt (C) expression (red in A-C and white in A’-C’) in stage-14 follicles with flip-out clones (marked by GFP) overexpressing *sim*^*RNAi*^ (*act>sim*^*RNAi*^). (D-F) Br-C (D), Cut (B), and Hnt (F) expression (red in D-F and white in D’-F’) in stage-14 follicles with flip-out clones (marked by GFP) overexpressing *tgo*^*RNAi1*^ (*act>tgo*^*RNAi1*^). The clone boundary is outlined by a blue dashed line. Nuclei are marked by DAPI in blue.

### The Sim:Tgo complex is required for ovulation

To further support that Sim and Tgo expression at stages 10-12 promotes follicle cell differentiation into maturation, we attempted to test whether knockdown of *sim* or *tgo* in follicle cells led to an ovulation defect as did knockdown of *ftz-f1* (Knapp et al., 2020). To do so, we used *Vm26Aa-Gal4*, a Gal4 driver expressed in follicle cells starting at stage 10 (Peters et al., 2013). Knockdown of *sim* or *tgo* using *Vm26Aa-Gal4* resulted in almost complete inhibition of egg laying (Fig 3A). In addition, we observed an increased number of stage-14 egg chambers inside the ovary, indicating an egg-retention phenotype and ovulation defect (Fig 3B). To determine whether the reduction in egg laying is indeed due to an ovulation defect, we estimated the time spent in ovulation using an assay described in previous reports (Deady and Sun, 2015; Sun and Spradling, 2013). *sim*- and *tgo*-knockdown females showed a dramatic increase in ovulation time. Control females usually take ∼5 min to ovulate an egg (Fig 3C and Table S1). In contrast, *sim*- and *tgo*-knockdown females take more than two hours to ovulate an egg (Fig 3C and Table S1).

**Figure 3.**
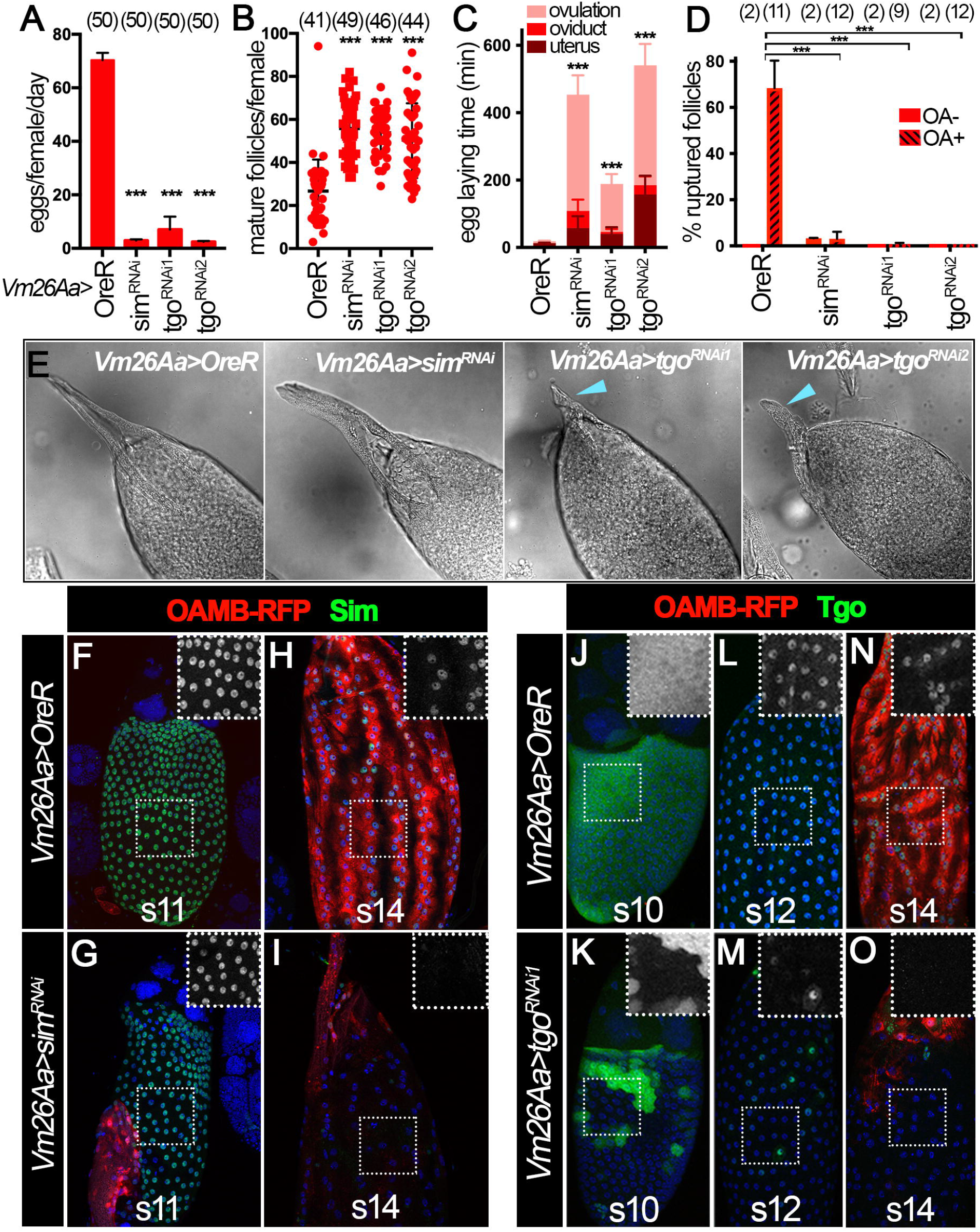
The Sim:Tgo complex is required for ovulation. **(A-C)** Quantification of egg laying (A), mature follicle numbers after egg laying (B) and egg laying time (C) in control, *sim*^*RNAi*^, *tgo*^*RNAi1*^ or *tgo*^*RNAi2*^ females with *Vm26Aa-Gal4* and *Oamb-RFP*. The number of females is noted above each bar. **(D)** Quantification of OA-induced follicle rupture using mature follicles isolated from control, *sim*^*RNAi*^, *tgo*^*RNAi1*^ or *tgo*^*RNAi2*^ females with *Vm26Aa-Gal4* and *Oamb-RFP*. The number of wells analyzed (containing 25-35 follicles) is noted above each bar. **(E)** Representative DIC images show dorsal appendages morphology in control, *sim*^*RNAi*^, *tgo*^*RNAi1*^ or *tgo*^*RNAi2*^ stage-14 follicles with *Vm26Aa-Gal4* and *Oamb-RFP*. Blue arrowheads indicate stunted dorsal appendages. **(F-I)** Sim expression (green) in stage-11 (F-G) and stage-14 follicles (H-I) from control (F, H) and *sim*^*RNAi*^ (G, I) females with *Vm26Aa-Gal4* and *Oamb-RFP*. The insets are higher magnification of Sim expression (white) in squared areas. **(J-O)** Tgo expression (green) in stage-10 (J-K), stage-12 (L-M), and stage-14 (N-O) follicles from control (J, L, N) and *tgo*^*RNAi1*^ (K, M, O) females with *Vm26Aa-Gal4* and *Oamb-RFP*. The insets are higher magnification of Tgo expression (white) in squared areas. Nuclei are marked by DAPI in blue. ***p<0.001 (Student’s t-test).

To further probe whether this ovulation defect is due to the follicle’s incompetency to undergo OA-induced follicle rupture, we attempted to isolate mature follicles and carry out an *ex vivo* OA-induced follicle rupture assay described previously (Deady and Sun, 2015; Knapp et al., 2018). The *Oamb-RFP* reporter was used as in previous experiments to identify the mature follicles with an intact follicle-cell layer. However, we immediately noticed that *sim*-knockdown follicles did not express OAMB-RFP in most of the follicle cells except anterior follicle cells (see description below). Therefore, we used anterior OAMB-RFP expression to identify the mature follicles and used the bright field signal to ensure these follicles had an intact follicle-cell layer (see MM). After a three-hour incubation with OA, we examined the follicle rupture by staining the tissue with DAPI. We found that all mature follicles isolated in this way had an intact follicle-cell layer before culture, and about 60% of follicles from control females ruptured after three hours (Fig 3D, Fig S2A). In contrast, follicles isolated from *sim*- and *tgo*-knockdown females showed less than 5% rupture rate (Fig 3D, Fig S2B-D).

During follicle isolation, we noticed that *tgo*-knockdown mature follicles were rounder with shortened dorsal appendages, reminiscent of *ftz-f1*-knockdown follicles (Knapp et al., 2020), while *sim*-knockdown follicles were morphologically normal (Fig 3E). This inconsistency prompted us to examine the knockdown efficiency in these genotypes. To our surprise, *sim*^*RNAi*^ driven by *Vm26Aa-Gal4* showed no obvious effect on Sim protein expression from stages 10B to 12 (Fig 3F-G), while both *tgo*^*RNAi*^ lines showed drastic reduction of Tgo protein from stages 10B to 12 (Fig 3J-M, S2E-F). This inefficient knockdown of Sim protein via *sim*^*RNAi*^ is further supported by the observation that in these knockdown follicles Tgo is properly localized to the follicle cell nuclei (Fig S2H-I) and that Sim::GFP fusion protein is still detected in these follicle cells (Fig S2K-N) from stages 10B to 12. At stage 14, both *sim*^*RNAi*^ and *tgo*^*RNAi*^ lines were efficient to knock down Sim and Tgo protein, respectively (Fig 3H-I, 3N-O, S2G, S2J, and S2O-P). In addition, both led to a severe reduction of OAMB-RFP expression.

To test whether the inefficient knockdown of Sim protein in *sim*^*RNAi*^ line is the cause for the morphological difference between *sim*^*RNAi*^ and *tgo*^*RNAi*^ follicles, we attempted to knock down *sim* using *C204-Gal4*, which starts to express in mainbody and posterior follicle cells at stage 8 (Manseau et al., 1997; Sun et al., 2008). Due to the limited Sim antibody, we used Tgo nuclear localization to infer the *sim* knockdown efficiency. We found that Tgo nuclear localization is significantly reduced in stages 10B-12 and 14 follicles with *C204-Gal4* driving *sim*^*RNAi*^ expression (Fig S3A-F), indicating that Sim protein is efficiently knocked down with this manipulation. Follicles with this manipulation showed shorter dorsal appendages and rounder morphology (reflected by a reduction in the follicles’ AP/DV ratio; Fig S3G-H), just like *tgo*- and *ftz-f1*-knockdown follicles (Fig 3E). In addition, follicle cell differentiation and maturation was disrupted, manifested by the extended expression of Br-C and Cut and the prevention of Hnt upregulation at stage 14 (Fig S3I-N). All these data support the conclusion that Sim and Tgo form a complex and function downstream of Ftz-f1 in promoting follicle cell differentiation and maturation, which is critical for successful ovulation.

### Re-upregulation of Sim at stage 14 is essential for ovulation

The fact that females with *Vm26Aa-Gal4* driving *sim*^*RNAi*^ expression showed a severe ovulation defect without affecting Sim expression at stages 10B-12 prompted us to propose that re-upregulation of Sim at stage 14 plays an active role in mediating follicle rupture and ovulation. To test this hypothesis explicitly, we used three stage-14 follicle cell Gal4 drivers to deplete Sim specifically in stage-14 follicle cells: *47A04-Gal4* (Deady and Sun, 2015), *44E10-Gal4* (Deady and Sun, 2015), and *CG13083-Gal4* (Knapp et al., 2019). All three Gal4 drivers start expression in stage-14 follicle cells with *CG13083-Gal4* at the earliest (possibly in late stage 13) and *47A04-Gal4* at the latest. We first examined the *sim* knockdown efficiency using these three Gal4 lines. Consistent with the Gal4 expression pattern, Sim protein is not reduced from stages 10B-12 in all three conditions (Fig S4A-B, D-E, G-H). In addition, Sim protein is not efficiently depleted in early stage-14 follicles (including stage 14A and 14B as defined in (Deady et al., 2017)) except in the case of *CG13083-Gal4*, which showed some reduction at this stage (Fig S4A-I). In contrast, Sim protein is clearly depleted in late stage-14 follicles with all three Gal4 drivers (Fig S4A-I). Next, we examined whether this manipulation caused an ovulation defect. Regardless of the Gal4 driver used, *sim* depletion with either of these Gal4 lines led to a significant decrease in egg laying (Fig 4A, S5A, S5J), mature follicle retention (Fig 4B, S5K), and an increase in ovulation time (Fig 4C, S5C, S5L, Table S1). All these data suggest that Sim in stage-14 follicle cells is essential for ovulation *in vivo*.

**Figure 4.**
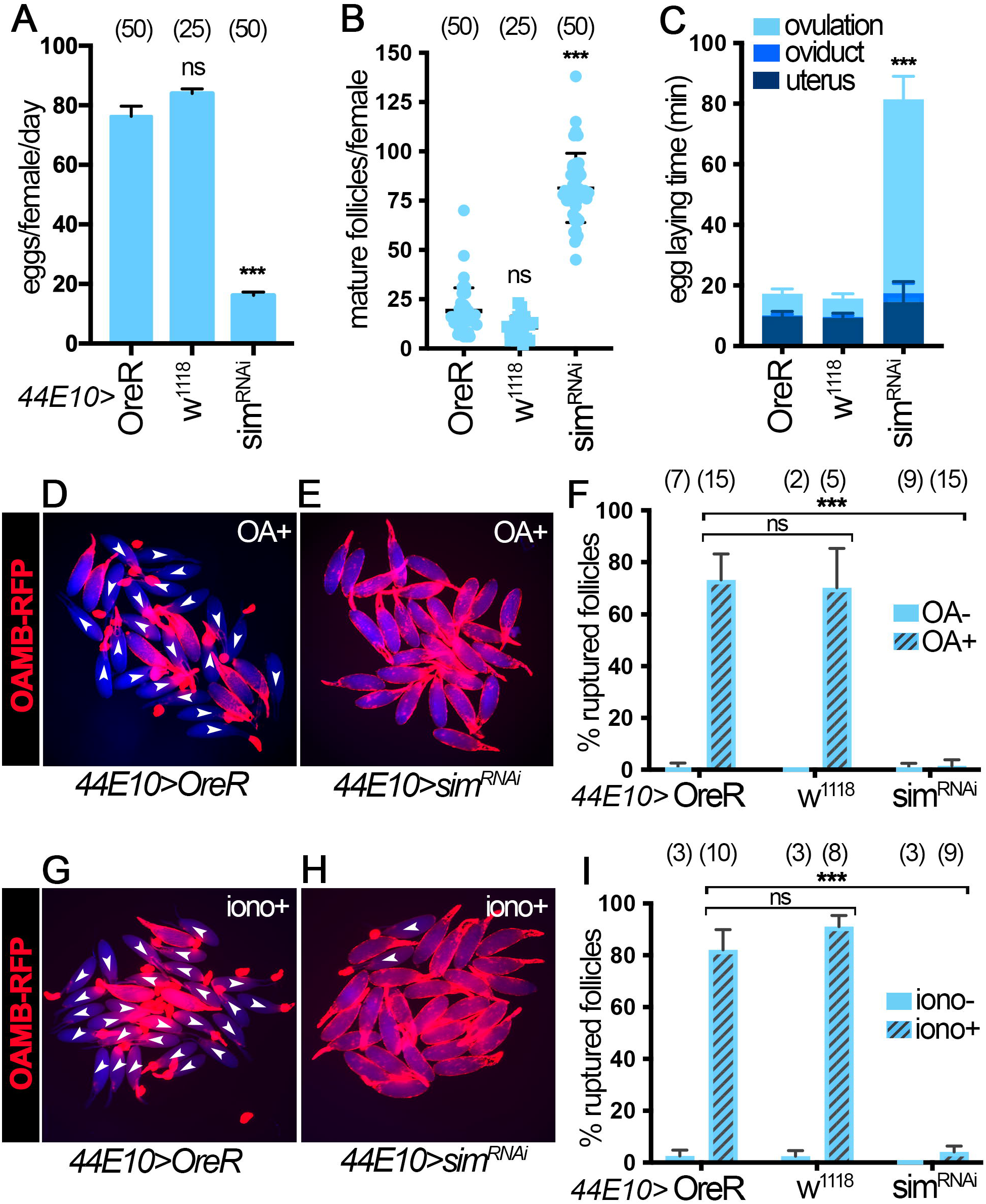
*Sim* knockdown with 44E10-Gal4 driver leads to ovulation defects. **(A-C)** Quantification of egg laying (A), mature follicle numbers after egg laying (B) and egg laying time (C) in control and *sim*^*RNAi*^ females with *44E10-Gal4* and *Oamb-RFP*. The number of females is noted above each bar. **(D-E**) Representative images show mature follicles from control (D) and *sim*^*RNAi*^ (E) females with *44E10-Gal4* and *Oamb-RFP* after 3h culture with 20μM OA (OA+). Oamb-RFP is shown in red and bright-field signal is shown in blue. Ruptured follicles are labeled with white arrowheads. (F) Quantification of OA-induced follicle rupture. The number of wells analyzed (containing 25-35 follicles) is noted above each bar. **(G-H)** Representative images show mature follicles from control (G) and *sim*^*RNAi*^ (H) females with *44E10-Gal4* and *Oamb-RFP* after 3h culture with 2 μM ionomycin (iono+). Oamb-RFP is shown in red and bright-field signal is shown in blue. Ruptured follicles are labeled with white arrowheads. (I) Quantification of ionomycin-induced follicle rupture. The number of wells analyzed (containing 25-35 follicles) is noted above each bar. ***p<0.001 (Student’s t-test).

Next, we set to determine whether *sim* is required for OA-induced follicle rupture using our *ex vivo* follicle rupture assay. Mature follicles from control females ruptured normally in response to OA treatment, whereas mature follicles from *sim*^*RNAi*^ females using *44E10-Gal4* or *47A04-Gal4* drivers showed a severe defect in follicle rupture (Fig 4D-F, Fig S5D-F). Due to the difficulty to isolate mature follicles with *CG13083-Gal4*-mediated knockdown of *sim*, which prevent OAMB-RFP expression, we did not perform the follicle rupture assay with this manipulation. In addition, we tested whether *sim*-depleted follicles can rupture in response to ionomycin, a Ca^2+^ ionophore, which can bypass OAMB to increase intracellular Ca^2+^ (Deady and Sun, 2015). Interestingly, *sim*-knockdown follicles exhibited significant follicle rupture defects in response to ionomycin stimulation (Fig 4G-I, S5G-I). All these data suggest that Sim in stage-14 follicle cells is required for follicle rupture and likely regulates components downstream of Ca^2+^, in addition to OAMB.

### Depletion of Sim using tunable CRISPR-Cas9 system results in similar defects as those observed in *sim*^*RNAi*^ females

The *sim*^*RNAi*^ used in this study has no predictable off targets. To further eliminate the off-target concern, we used a tunable CRISPR-Cas9 system (Port et al., 2020) to disrupt Sim expression. The ability of the CRISPR-Cas9 system to efficiently knock out genes in follicle cells in late oogenesis has never been tested, since follicle cells undergo three rounds of endoreplication and contain 16 copies of the genome. To this end, we used follicle cell-specific *Vm26Aa-Gal4* driving two guide RNAs targeting the first and second common coding exons of the *sim* gene, as well as different levels of Cas9 expression in follicle cells. *UAS-Cas9*^*XS*^ expresses Cas9 at the highest levels, and *UAS-Cas9*^*M*^ expresses Cas9 at the lowest levels (Port et al., 2020). This genetic manipulation disrupts Tgo nuclear localization (inferring Sim expression) in stage-14 follicle cells but not before stage 14 (Fig 5A-D). In addition, these females showed reduced egg laying capacity and increased mature follicle number in ovaries (Fig 5E-F), reminiscent of *sim*-knockdown females. Most importantly, mature follicles from *sim*-knockout females with the CRISPR-Cas9 system also showed defective OA-induced follicle rupture and reduced OAMB-RFP expression (Fig 5G-I). These data demonstrate that tissue-specific CRISPR-Cas9 system is sufficient to knock out genes in follicle cells and that Sim in stage-14 follicle cells play an active role in mediating follicle rupture and ovulation.

**Figure 5.**
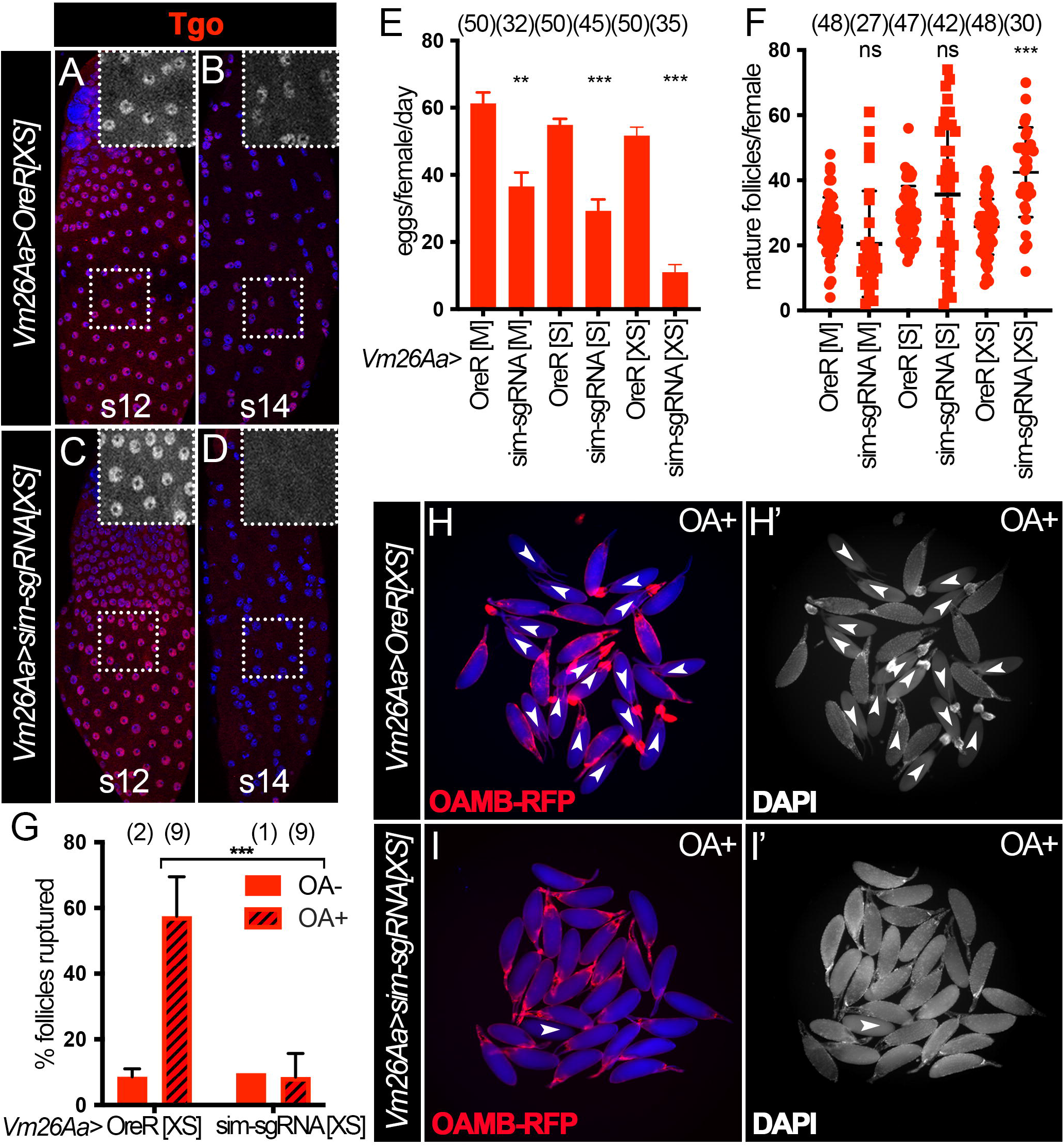
Depletion of Sim using tunable CRISPR-Cas9 system results in similar defects as those in *sim*^*RNAi*^ females. **(A–D)** Tgo expression (red) in stage-12 (A, C) and stage-14 (B, D) follicles from control (A-B) and *sim*^*sgRNA*^ (C-D) females with *Vm26Aa-Gal4* and *UAS-Cas9*^*XS*^. The insets are higher magnification of Tgo expression (white) in outlined areas. All images from A-D were acquired using the same microscopic settings. Nuclei are marked by DAPI in blue. **(E-F)** Quantification of egg laying (E) and mature follicle retention (F) in control and *sim*^*sgRNA*^ females with *Vm26Aa-Gal4* and three different *UAS-Cas9* constructs: M, S and XS. The number of females is noted above each bar. (**G**) Quantification of OA-induced follicle rupture assessed via DAPI staining. The number of wells analyzed is noted above each bar. (**H-I**) Representative images show mature follicles from control (H) and *sim*^*sgRNA*^ (I) females with *Vm26Aa-Gal4, UAS-Cas9*^*XS*^, and *Oamb-RFP* after 3h culture with 20 μM OA (OA+). Oamb-RFP is shown in red and bright-field signal is shown in blue, and DAPI signal is shown in white in (H’-I’). Ruptured follicles are labeled with white arrowheads and usually have corpus luteum accumulation at the posterior end by the dorsal appendage. ***p<0.001 (Student’s t-test).

### Sim at stage 14 upregulates *Oamb* expression in mature follicles in conjunction with Hnt

To explore the mechanism of Sim in ovulation, we investigated whether Sim regulates known ovulatory genes. As mentioned previously, mature follicles with *Vm26Aa-Gal4* driving *sim*^*RNAi*^ expression showed almost no detectable *Oamb-RFP* in most of the follicle cells (Fig 3I).We hypothesized that Sim is required for *Oamb* transcription in mature follicle cells. Consistent with this hypothesis, mature follicles with *CG13083-Gal4* driving *sim*^*RNAi*^ expression also showed no detectable *Oamb-RFP* in most follicle cells except in the anterior region (Fig 6A-B, F, S6A-D). In contrast, knockdown of *sim* with *44E10-Gal4* or *47A04-Gal4* had much weaker (if any) effect on *Oamb-RFP* expression (Fig S6E-L) but still showed significant reduction of *Oamb* mRNA level (Fig S6M-N). All these data indicate that Sim in stage-14 follicles is required for *Oamb* transcription.

**Figure 6.**
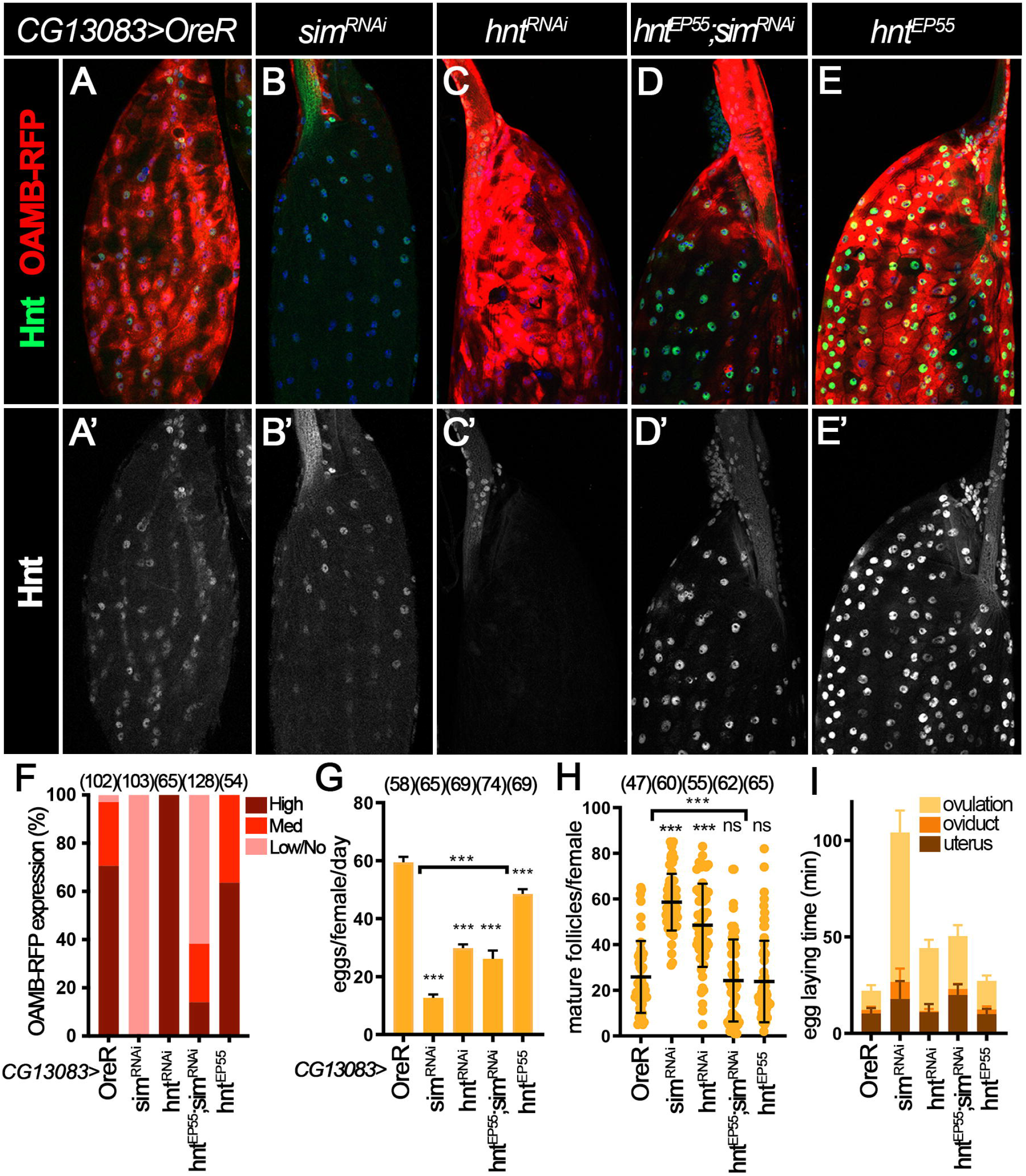
Sim at stage 14 upregulates *Oamb* expression in mature follicles in conjunction with Hnt. **(A-E)** Hnt expression (green in A-E and white in A’-E’) and OAMB-RFP expression (red in A-E; detected by RFP antibody) in control(A), *sim*^*RNAi*^ (B), *hnt*^*RNAi*^ (C), *hnt*^*EP55*^*;sim*^*RNAi*^ (D), and *hnt*^*EP55*^ (E) stage-14 follicles with *CG13083-Gal4 and Oamb-RFP*. Nuclei are marked by DAPI in blue. **(F)** Quantification of main body OAMB-RFP expression levels in late stage-14 follicles (characterized by absence of nurse cell nuclei, follicle morphology and high RFP expression in the dorsal appendages region). The number of mature follicles quantified is noted above each bar. **(G-I)** Quantification of egg laying **(G)**, mature follicles retained **(H)** and egg laying time **(I)** in control, *sim*^*RNAi*^, *hnt*^*RNAi*^, *hnt*^*EP55*^*;sim*^*RNAi*^ and *hnt*^*EP55*^ females with *CG13083-Gal4* and *Oamb-RFP*. The number of females is noted above each bar. ***p<0.001 (Student’s t-test).

Our previous work showed that Hnt upregulation in stage-14 follicle cells is required for *Oamb* transcription (Deady et al., 2017). We therefore examined Hnt expression in *sim*-knockdown follicles with *CG13083-Gal4* driver. We observed that upon *sim* knockdown, Hnt could still be upregulated in early and mid stage-14 follicle cells (Fig S6A and S6C) but showed an obvious reduction in late stage-14 follicle cells (Fig 6A-B, S6B and S6D). A similar reduction of Hnt expression in late stage-14 follicle cells was also observed when *sim*^*RNAi*^ was driven by *44E10-Gal4* (Fig S6F and S6H) but this reduction was not obvious with *47A04-Gal4* (Fig S6J and S6L). Consistent with the protein expression, *hnt* mRNA is also drastically reduced in follicle cells with *44E10-Gal4* but not *47A04-Gal4* driving *sim*^*RNAi*^ expression (Fig S6M-N). Together, these data indicate that Sim in stage-14 follicles also regulates Hnt expression.

To test whether Sim activates *Oamb* expression via Hnt, we examined whether *hnt* overexpression (with *UAS-hnt*^*EP55*^) in *sim*-knockdown follicles could restore *Oamb-RFP* expression. Using this genetic manipulation, Hnt protein is restored in mature follicle cells (Fig 6D). Surprisingly, the defective *Oamb-RFP* expression associated with *sim*-knockdown could not be fully rescued by *hnt* overexpression (Fig 6D-F). In addition, *Oamb* mRNA levels were also not rescued by *hnt* overexpression in *sim*-knockdown follicles with *47A04-Gal4* (Fig S6N). These results indicate that *hnt* is not the only target downstream of Sim to regulate *Oamb* transcription. Consistent with this idea, *hnt*-knockdown using *CG13083-Gal4* did not affect *Oamb-RFP* expression, unlike *sim*-knockdown follicles (Fig 6C). In addition, *hnt* overexpression only showed a partial rescue of egg laying and ovulation defects in *sim*-knockdown follicles (Fig 6G-I, Table S1). Therefore, these results lead us to conclude that Sim regulates OAMB expression in mature follicle cells independent of and in conjunction with Hnt.

### Sim at stage 14 upregulates Mmp2 in mature follicles in conjunction with Hnt

The fact that ionomycin failed to induce follicle rupture in *sim*-knockdown follicles (Fig 4G-I and S5G-I) suggests that *sim* could regulate components downstream of Ca^2+^, such as Mmp2 and ROS signaling components. To test whether late Sim regulates Mmp2 expression in posterior follicle cells, we examined *Mmp2::GFP* reporter expression in *sim*-knockdown follicles. Due to technical difficulties to recombine *CG13083-Gal4* and *Mmp2::GFP*, we were unable to examine *Mmp2::GFP* expression in *sim*-knockdown follicles with *CG13083-Gal4*.Instead, we examined *Mmp2::GFP* expression in *sim*-knockdown follicles with *44E10-Gal4* or *47A04-Gal4*. We found that Mmp2::GFP is significantly reduced in posterior follicle cells of late stage-14 follicles with *sim* knockdown relative to controls (Fig 7A-B, E-G). Moreover, *Mmp2* mRNA levels were also significantly reduced in *sim*-knockdown follicles (Fig S6M-N). All these data indicate that Sim is required for Mmp2 expression.

**Figure 7.**
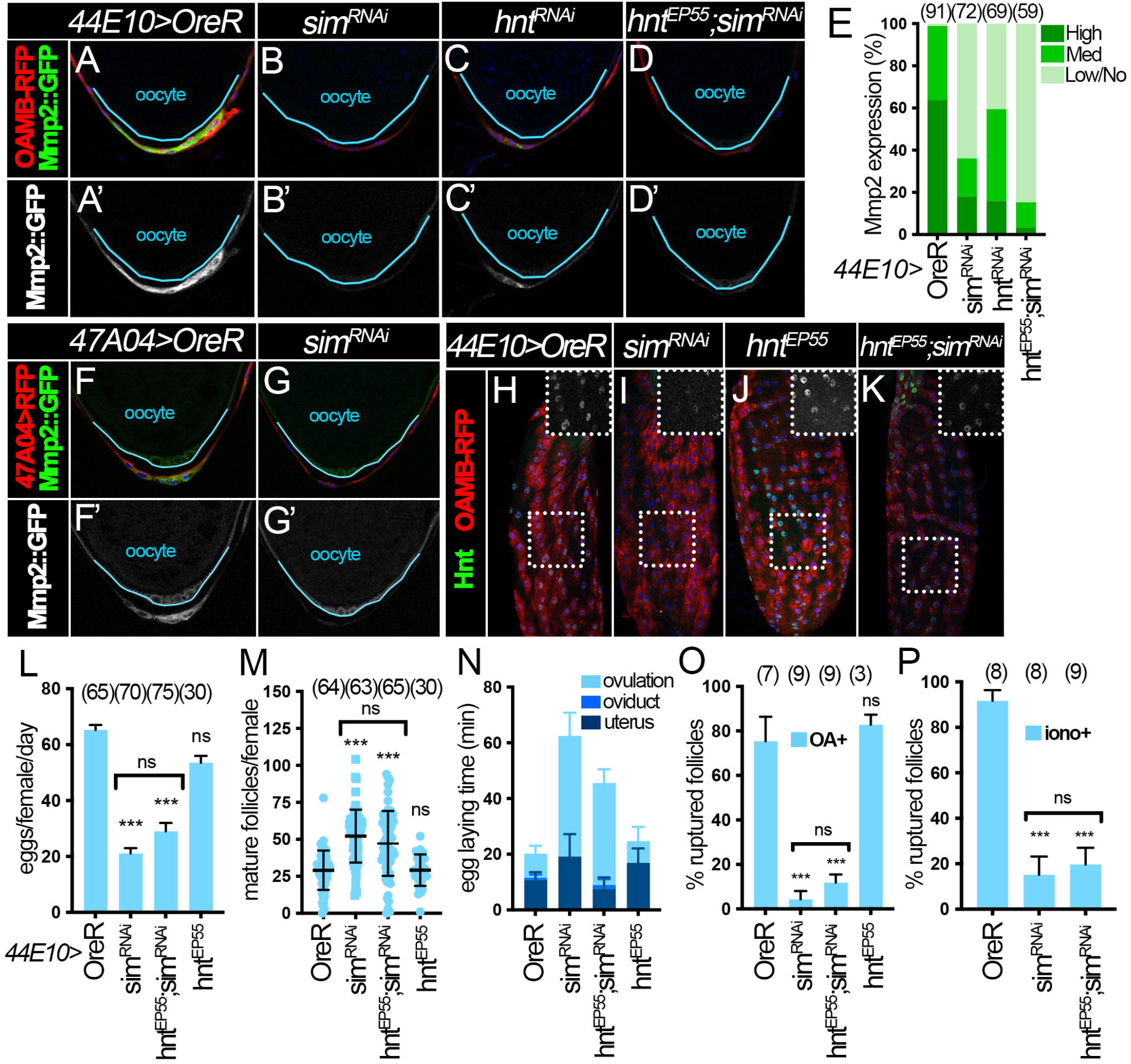
Sim at stage 14 upregulates Mmp2 in mature follicles in conjunction with Hnt. **(A-D)** Representative images show Mmp2::GFP expression (green in A-D and white in A’-D’) in posterior follicle cells of control (A), *sim*^*RNAi*^ (B), *hnt*^*RNAi*^ (C), and *hnt*^*EP55*^*;sim*^*RNAi*^ (D) mature follicles with *44E10-Gal4* and *OAMB-RFP*. Oocytes are outlined in blue. All images from A-D were acquired using the same microscopic settings. Nuclei are marked by DAPI in blue in all panels. **(E)** Quantification of Mmp2::GFP expression levels in posterior follicle cells of unruptured late stage-14 follicles (characterized by highest RFP expression, long dorsal appendages and absence of nurse cell nuclei). Posterior polar cell expression is not considered when quantifying Mmp2::GFP expression intensity. The number of mature follicles quantified is noted above each bar. **(F–G)** Representative images show Mmp2::GFP expression (green in F-G and white in F’-G’) in control (F) and *sim*^*RNAi*^ (G) late stage-14 follicles marked by *47A04-Gal4*>*UAS-RFP* (red). Oocytes are outlined in blue. All images from F-G were acquired using the same microscopic settings. **(H–K)** Hnt expression (green) in late stage-14 follicles from control (H), *sim*^*RNAi*^ (I), *hnt*^*EP55*^ (J), and *hnt*^*EP55*^*;sim*^*RNAi*^ (K) females with *44E10-Gal4* and *Oamb-RFP*. Late stage-14 follicle cells are marked by the highest level of OAMB-RFP expression (red). The insets are higher magnification of Hnt expression (white) in outlined areas. All images from H-K were acquired using the same microscopic settings. **(L-N)** Quantification of egg laying (L), mature follicles retained (M) and egg laying time (N) in control, *sim*^*RNAi*^, *hnt*^*EP55*^*;sim*^*RNAi*^ and *hnt*^*EP55*^ females with *44E10-Gal4* and *Oamb-RFP*. The number of females is noted above each bar. **(O-P)** Quantification of OA-induced follicle rupture (O) and ionomycin-induced follicle rupture (P) in control, *sim*^*RNAi*^, *hnt*^*EP55*^*;sim*^*RNAi*^ and *hnt*^*EP55*^ females with *44E10-Gal4* and *Oamb-RFP*. The number of wells analyzed (containing 25-35 follicles) is noted above each bar. ***p<0.001; ns: Not significant (Student’s t-test).

We also noticed that *sim-*knockdown follicles showed a more severe Mmp2 reduction than *hnt*-knockdown follicles (Fig 7C and E), which prompted us to investigate whether Sim regulates Mmp2 expression via Hnt. We evaluated Mmp2::GFP expression in follicle cells with *sim* knockdown and *hnt* overexpression simultaneously, and found that these follicles showed similar (if not worse) Mmp2::GFP reduction as *sim*-knockdown follicles (Fig 7D-E, H-K). In addition, *hnt* overexpression did not rescue the ovulation and follicle rupture defect caused by *sim* knockdown (Fig 7L-P, Table S1), similar to the results with *CG13083-Gal4* (Fig 6G-I). Altogether, our results suggest that Sim regulates Mmp2 expression independent of and in conjunction with Hnt.

### Sim is required for NOX expression independent of Hnt

Next, we examined whether Sim regulates NOX expression in stage-14 follicles. Our previous work indicates that NOX is present in stage-14 follicle cells to produce ROS required for follicle rupture (Li et al., 2018). However, NOX expression throughout oogenesis and its regulation is completely unknown. To address this question, we generated *Nox::GFP* knock-in allele by replacing *Nox*^*CRIMIC*.*TG4*.*0*^ allele with the Double Header *GFP/T2A-Gal4* coding cassette (Li-Kroeger et al., 2018). NOX::GFP was not detected in follicle cells before stage 13, and it was faintly detected in a few follicle cells at late stage 13 (Fig 8A). Its expression started to upregulate at early stage 14 and peaked at late stage 14 (Fig 8B-C). This upregulation of NOX::GFP is earlier than the upregulation of Hnt (Fig 8B), suggesting that Hnt may not regulate NOX expression. Consistent with this idea, *hnt* knockdown with *CG13083-Gal4* did not disrupt NOX::GFP expression in mature follicle cells (Fig 8D). In contrast, NOX::GFP was drastically reduced in *sim*-knockdown follicles with *CG13083-Gal4* (Fig 8E). Consistent with the NOX::GFP protein expression defect, OA- and ionomycin-induced superoxide production were significantly reduced in *sim*-knockdown follicles (Fig 8I-J). It is interesting to note that *sim* knockdown with *44E10-Gal4* or *47A04-Gal4* did not show a clear reduction of NOX::GFP expression (Fig S7A-D) but showed the reduction of *Nox* mRNA (Fig S6M-N). In addition, *hnt* overexpression did not restore NOX::GFP expression in *sim*-knockdown follicles with *CG13083-Gal4* (Fig 8F-G), nor did it restore *Nox* mRNA expression in sim-knockdown follicles with *47A04-Gal4* (Fig S6N). All these data suggest that Sim upregulates NOX expression in stage-14 follicle cells independent of Hnt. In conclusion, we demonstrated that Sim functions like a master regulator in late follicles to regulate ovulatory competency by upregulating all known ovulatory genes, including OAMB, Mmp2, and NOX (Fig 8K).

**Figure 8.**
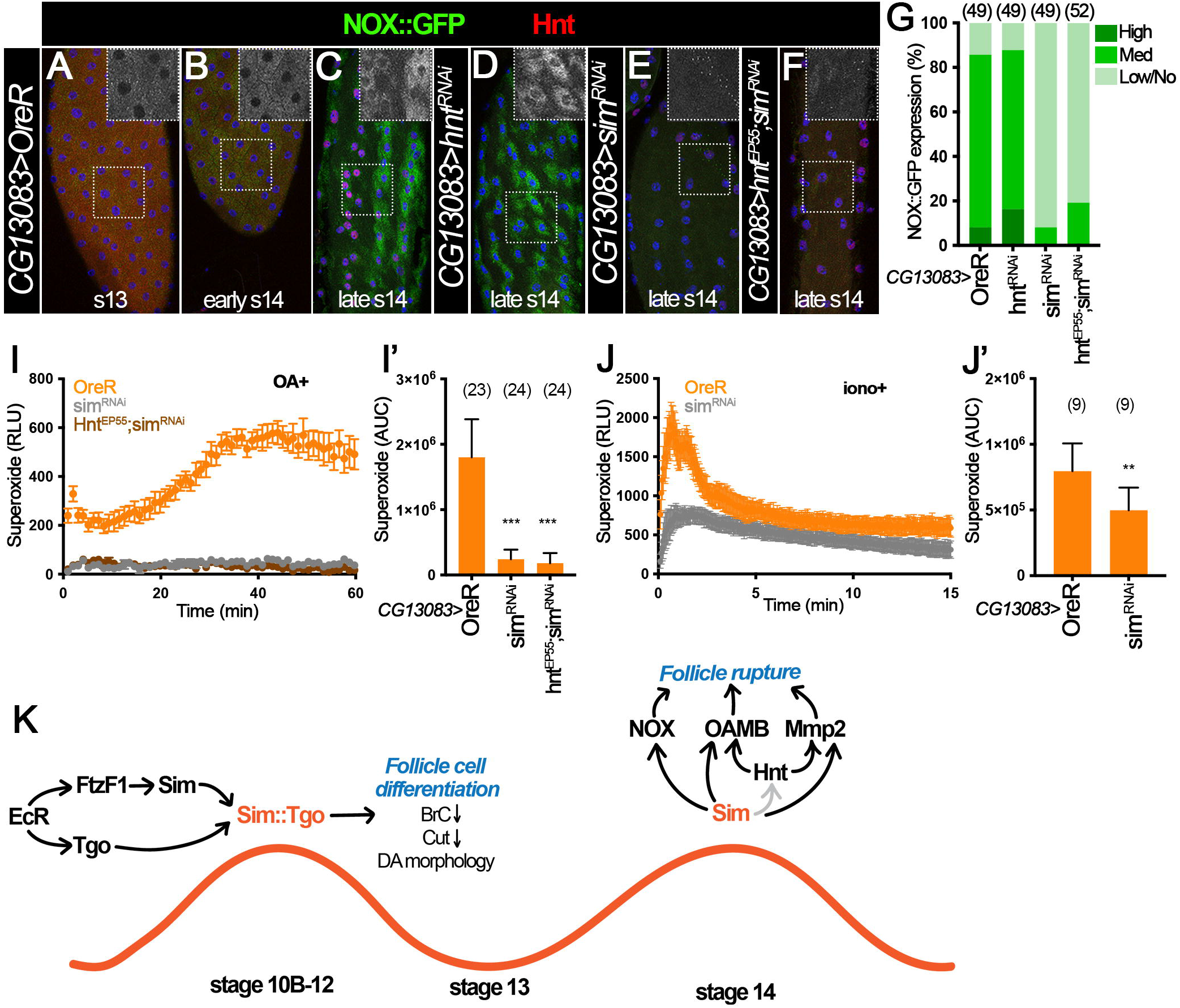
Sim is required for NOX expression independent of Hnt. **(A–C)** NOX::GFP (green, detected by anti-GFP antibody) and Hnt (red) in stage-13 (A), early stage-14 (B) and late stage-14 (C) follicles from *NOX::GFP* control females with *CG13083-Gal4*. The insets are higher magnification of NOX::GFP expression (white) in squared areas. **(D-F)** NOX::GFP (green) and Hnt expression (red) in *hnt*^*RNAi*^ (D), *sim*^*RNAi*^ (E), and *hnt*^*EP55*^*;sim*^*RNAi*^ (F) late stage-14 follicles with *CG13083-Gal4*. All images from A-F were acquired using the same microscopic settings. Nuclei are marked by DAPI in blue. (G) Quantification of NOX::GFP expression levels in late stage-14 follicles (characterized by long dorsal appendages, lack of nurse cell nuclei and Hnt expression). The number of mature follicles quantified is noted above each bar. (**I-J**) Measurement of OA-induced (I) and ionomycin-induced (J) superoxide production in stage-14 follicles from control (orange), *sim*^*RNAi*^ (gray), and *hnt*^*EP55*^*;sim*^*RNAi*^ (brown) females with *CG13083-Gal4* and *Oamb-RFP*. The superoxide is detected by L-012 luminescent signal (relative luminescence). **(I’-J’)** Quantification of the area under the curve representing superoxide production for each genotype and condition. The number of wells analyzed is noted above each bar. **p<0.01 ***p<0.001 (Student’s t-test). **(K)** A cartoon illustrates the role of Sim and Tgo in follicle differentiation and ovulation in late oogenesis in Drosophila. Downstream of ecdysteroid signaling, Sim and Tgo form a complex to promote follicle differentiation by downregulating Br-C and Cut and regulating follicle morphology at stage10B-12. Sim is re-upregulated at stage 14 and plays an active role in regulating the expression of Hnt, Mmp2, NOX and OAMB proteins required for OA-induced follicle rupture.

## Discussion

Development of a mature follicle is a long and complex process conserved across metazoans and requires the orchestration of many signaling pathways between somatic follicle cells and germ cells to ensure their proper differentiation and maturation (Merkle et al., 2020; Doherty et al., 2022). A large body of work has focused on the formation and development of follicles during early stages (Duhart et al., 2017; Spradling et al., 2022), while less work has focused on the late follicle development and the maturation process that leads to their competency for ovulatory stimuli. Our previous work in *Drosophila* identified the nuclear receptor Ftz-f1, which plays a conserved role in follicle differentiation and maturation during late oogenesis downstream of ecdysteroid signaling (Knapp et al., 2020). Ftz-f1 directly activates the bHLH-PAS transcription factor Sim. However, the mechanism of Sim in follicle differentiation and maturation is unknown. This paper contributes to this knowledge gap by uncovering how the bHLH-PAS protein complex Sim:Tgo functions within the *Drosophila* ovulatory network to regulate key genes required for the follicle maturation and follicle rupture pathways, which are conserved in preovulatory mammalian follicles to ensure reproductive success.

We discovered that ecdysteroid signaling at stage10A/10B transition not only activates Ftz-f1, but also upregulates another bHLH-PAS protein Tgo (Fig 8K), the heterodimeric partner of Sim (Ohshiro and Saigo, 1997; Sonnenfeld et al., 1997). Sim and Tgo then form a complex, translocate into the nucleus, and promote follicle cell differentiation, including downregulating Br-C and Cut. Sim:Tgo represses gene expression by indirectly activating the repressive factors in CNS development (Estes et al., 2001). Therefore, we suspect that downregulation of Br-C and Cut by Sim:Tgo may also be indirect. This is also consistent with the fact that downregulation of Br-C and Cut is observed at stage 13 when Sim and Tgo are also downregulated (Fig 1D). Future work will focus on identifying the molecular mechanism of this regulation.

Sim:Tgo’s role in follicle maturation does not seem to be limited to stages 10B to 12 as both Sim and Tgo are re-upregulated in stage-14 follicle cells after a transient downregulation at stage 13 (Fig 1). With multiple stage-14 Gal4 drivers, we were able to disrupt late Sim expression (stage 14) without altering its early expression (stages 10B-12). Our data strongly suggests that late Sim acts as a master regulator of all known genes involved in follicle rupture and ovulation, including *hnt, Nox, Oamb*, and *Mmp2* (Fig 8K). Unfortunately, we were unable to efficiently disrupt late Tgo expression using stage-14 Gal4 drivers (data not shown). Therefore, we were unable to provide conclusive evidence that Tgo also partners with Sim to regulate ovulatory gene expression in stage-14 follicle cells. However, Tgo’s nuclear localization in stage-14 follicle cells also depends on Sim (Fig S2J), and Sim and Tgo form a complex to regulate gene expression in every studied tissue or cell type to date (Sonnenfeld et al., 1997; Estes et al., 2001). Consequently, we suspect that Tgo is still the partner of Sim in stage-14 follicle cells. It is worth noting that it only takes less than 10 hours for follicles to develop from stages 10B to 14 (Spradling, 1993). Therefore, we find it intriguing to see multiple genes turned on and off in this short period of time, which suggests that the regulation of follicle maturation is highly complex and dynamic. A comprehensive understanding of this process in flies and other model systems may allow us to identify genetic causes for ovulation-related infertility, including polycystic ovarian syndrome, and lead to better treatment development for these health problems.

We also identified, for the first time, a factor that regulates NOX expression in follicle cells. Our work showed that late Sim is critical to upregulates *Nox* expression in follicle cells (Fig 8A-G). It is unclear whether Nox is a direct target of Sim. Previous work in CNS midline development identified Sim binding motif as ACGTG (Wharton et al., 1994). Further work by Hong and his group also found that Sim:Tgo could bind to a slight variable motif DDRCGTG (Hong et al., 2013; Shin and Hong, 2015; Shin and Hong, 2016). Using the latter motif to search the *Nox* gene region, we found six potential Sim binding motifs within 3kb upstream of *Nox-RC* isoform, which is the isoform expressed in follicle cells according to our RNAseq experiment (Knapp et al., 2020). In addition, there are 14 additional DDRCGTG motifs in the *Nox* gene region. Therefore, Sim could potentially regulate *Nox* expression through direct binding to its enhancer. In addition to *Nox*, Sim may directly regulate *Mmp2* and *Oamb* expression. There are three DDRCGTG motifs clustered within 350bp in the Mmp2 enhancer that directs Mmp2 expression in follicle cells (data presented in a different manuscript). There are two motifs clustered together in the *Oamb-RFP* enhancer. Our data showed that Hnt does not seem to regulate the *Oamb-RFP* enhancer (Fig 6C) despite of the fact that it can regulate *Oamb* gene expression (Deady et al., 2017). This information suggests that Hnt and Sim likely bind to different enhancer regions of *Oamb* gene if both Hnt and Sim regulate *Oamb* expression directly. Future work will be required to decipher whether Sim directly regulates the expression of these genes. Moreover, it would be interesting to identify additional differentially up-or down-regulated genes in *sim*-knockdown follicles using RNASeq, with the hope of finding other novel genes that could play critical roles in follicle maturation and ovulation that are unknown to date. These discoveries will aid in the process of understanding the precise transcription factor network that is spatiotemporally controlling follicle maturation and ovulation, key processes for reproductive success.

Sim:Tgo’s role in follicle maturation and ovulation may be conserved across metazoans. Both Ftz-f1 and its mammalian homology SF-1 play an important role in promoting follicle development and maturation (Knapp et al, 2020; Pelusi et al., 2008). In addition, many of the ovulatory genes and signaling pathways downstream of Sim also play conserved roles in *Drosophila* and mammalian ovulation, including matrix metalloproteinases, the adrenergic system, and ROS (Curry and Smith, 2006; Deady and Sun, 2015; Deady et al., 2015; Kannisto et al., 1985; Li et al., 2018; Shkolnik et al., 2011).

There are two mammalian *sim* homologs (*sim1* and *sim2*) (Chrast et al., 1997; Ema et al., 1996; Kewley et al., 2004).These mammalian Sim proteins also require heterodimerization with their partner protein ARNT, which is homologous to Tgo (Ema et al., 1996; Michaud et al., 2000). Interestingly, both *Drosophila* and mammalian Sim (murine, specifically) have been shown to play important roles in CNS development (Michaud et al., 1998; Michaud et al., 2000).Published RNAseq data evaluating human granulosa cells (Yerushalmi et al., 2014) has demonstrated that hSIM2 is induced after ovulation, whereas hSIM1 is not detected. Data reported in the Human Protein Atlas (Uhlén et al., 2015) also reports that hSIM2 is expressed at medium levels in follicle cells, whereas hSIM1 is not expressed in the human ovary. Together, this information suggests that it is highly likely that Sim and its mammalian partner ARNT play a role in mammalian follicle maturation.

## Materials & Methods

### Drosophila genetics

*Drosophila melanogaster* flies were reared on standard cornmeal-molasses food at 25°C except otherwise noted. All RNAi-mediated depletion experiments included the *UAS-dcr2* and were carried out at 29°C upon animal eclosion to increase knockdown efficiency. The following Gal4 drivers were used: *47A04-Gal4* and *44E10-Gal4* from the Janelia Gal4 collection (Pfeiffer et al., 2008), *CG13083-Gal4* (Knapp et al., 2019), *Vm26Aa-Gal4* (Peters et al., 2013), and *C204-Gal4* (Manseau et al., 1997; Sun et al., 2008). The *C204-Gal4* was the only Gal4 line that also contained *tub-Gal80*^*ts*^ to avoid early developmental defects. To knockdown or overexpress genes under the control of the above mentioned Gal4 drivers, the following transgenic lines were used: *UAS-sim*^*RNAi*^ (Vienna *Drosophila* Resource Center - VDRC, stock #26888), *UAS-tgo*^*RNAi1*^ (Bloomington *Drosophila* Stock Center - BDSC, stock #53351), *UAS-tgo*^*RNAi2*^ (BDSC, stock #26740), *UAS-hnt*^*RNAi*^ (VDRC, stock #3788), *UAS-hnt*^*EP55*^ (BDSC, stock #5358), and *UAS-EcR*^*DN*^ (BDSC, stock #6872). Isolation and identification of stage-14 follicles for follicle rupture assay were performed using *Oamb-RFP* (Knapp et al., 2019) or *47A04-Gal4* driving *UAS-RFP* (*47A04>RFP*) *or UAS-RG6* (47A04>RG6; Jiang et al., 2021). The following protein trap lines were used: *Mmp2::GFP* (Deady et al., 2015), *Sim::GFP* (VDRC, stock #318096; (Sarov et al., 2016), and *NOX::GFP*. The *Nox::GFP* reporter is generated by replacing *Nox*^*CRIMIC*.*TG4*.*0*^ allele with the Double Header GFP/T2A-Gal4 coding cassette following the exact cross scheme in (Li-Kroeger et al., 2018). For the CRISPR-Cas9 mediated disruption, *Vm26Aa-Gal4* was used to drive tunable *UAS-Cas9* constructs (XS, S, M; VDRC stock #340000, #340001, and #340002, respectively; (Port et al., 2020) and *UAS-sim*^*sgRNA*^ (VDRC, stock #341711). Control flies for all experiments were prepared by crossing Gal4 driver to *Oregon-R* or *w*^*1118*^ flies.

### Clone induction and generation of mosaics

For generation of flip-out clones, the *hsFLP;; act<CD2<Gal4, UAS-GFP/TM3* stock was crossed to the indicated transgenes of interest. For clone induction, adult female progeny with correct genotypes were heat shocked for 45 min at 37°C to induce FLP/FRT mediated recombination and incubated at 25°C for 2-4 days before dissection. During the time post-heat shock, flies were mated and provided with wet yeast paste accordingly to induce the production of the desired follicle stages required for the experiment (Jia et al., 2016).

### Ovulation and follicle rupture assays

The egg laying and mature follicle retention assays were conducted as described in previous literature, along with the egg laying time analysis (Deady and Sun, 2015; Knapp and Sun, 2017). In short, five-to six-day old virgin females fed with wet yeast for 1-day were used to access the number of eggs laid on a two-day period after mating with Oregon-R males, or to examine the location of eggs in the reproductive tract after six-hour mating with Oregon-R males. These data were then used to calculate the average number of eggs laid/female/day and the average time of ovulating an egg.

The OA and ionomycin-induced follicle rupture assays were performed as previously described (Knapp et al., 2018) except for the following modification when using *Vm26Aa-Gal4* driver. Due to the loss of Oamb-RFP reporter in sim-knockdown follicle cells, we were unable to use Oamb-RFP to isolate mature follicles and examine follicle rupture as in previous work.Instead, we relied on high levels of OAMB-RFP in anterior follicle cells to identify the mature follicles and used the bright field to ensure these follicles had an intact follicle-cell layer at the posterior (a transparent layer of follicle cells over the oocyte’s white yolk at 80x magnification). A couple of groups of 25-35 follicles isolated using this method were fixed and DAPI stained in a master fixation buffer (4% EM-grade paraformaldehyde + 0.1μg/mL DAPI) to determine that these follicles indeed had an intact layer of follicle cells. The rest of mature follicles in groups of 25-35 were incubated for 3h with OA medium (Grace’s Medium + 10% FBS + 100 U/ml penicillin-streptomycin + 20 μM OA) (Knapp et al., 2018). After incubation, follicles were fixed and DAPI stained in the master fixation buffer for 15 min. In all instances, washing was not performed after fixation/DAPI staining to avoid washing away the corpus luteum and disturbing the follicles. The follicles were immediately visualized and imaged using a Leica MZ10F fluorescence stereoscope, and the number of ruptured follicles was quantified according to DAPI signal (marking follicle cell nuclei). If more than 75% of the follicle lacked DAPI signals and an accumulation of follicle cells expressing DAPI is observed anteriorly, we count that follicle as ruptured, similar to what was previously described (Knapp et al., 2018). Each data point represents the percentage (mean percentage ± standard deviation (SD)) of ruptured follicles per group of ∼25-35 follicles.

### qRT-PCR

For quantitative RT-PCR, total RNA was extracted from 60 stage-14 follicles using TRIzol (Invitrogen) and the Direct-zol RNA MicroPrep Kit (Zymo Research, Orange, CA). Fifteen five-to six-day old virgin flies were fed wet yeast 3-days prior to dissection. Mature follicles were isolated using the *Oamb-RFP* reporter or the *47A04>RFP*. cDNA synthesis and real-time PCR amplification with three technical repeats were performed as previously described (Knapp and Sun, 2017; Li et al., 2018). Primers for *oamb*.*K3, mmp2* and *nox* were described previously (Knapp and Sun, 2017; Li et al., 2018). The *hnt* primer pair used was 5′-ACATCCGGTGCCACAATTAC-3′ and 5′-GTGAACGTCAGGTGGCAGTAG-3′. The *Rps17* primer was used as an internal control. The data are presented as mean ± SD from at least two biological replicates, to ensure reproducibility.

### Superoxide detection

Measurement of superoxide production was performed as previously described (Li et al., 2018) with slight modifications. Five-to six-day old virgin flies were fed wet yeast 3-days prior to dissection. Five intact mature follicles were isolated using *OAMB-RFP* and bright field and placed in each well of a 96-well plate injected with 100 μl of Grace’s insect medium containing 200 μM of L-012 (Wako Chemicals) and either 20 μM OA or 2 μM ionomycin. For OA-induced ROS detection, plates were placed in a CLARIOstar microplate reader (BMG Labtech) for luminescence reading for 60 min (60 cycles, 63 seconds each, measurement interval time 0.5 seconds). For ionomycin-induced ROS detection, luminescence reading lasted 15 min (300 cycles, 3 seconds each, measurement interval time 0.5 seconds, which was enough to catch the fast initial peak. A minimum of 3 wells (technical repeats) were used in each experiment per genotype, and the mean ± SEM of the technical repeats was calculated. Each experiment was performed at least twice.

### Immunostaining and microscopy

Adult female ovaries were dissected in cold Grace’s medium and fixed in 4% EM-grade paraformaldehyde in PBT (1XPBS + 0.2% Triton X-100) for 13-15 min at room temperature.The tissue was washed vigorously with PBT, blocked in antibody buffer (PBT + 0.5% BSA+ 2% normal goat serum), and stained with primary antibodies overnight at 4°C. After primaries, tissue was washed with PBT and incubated in secondary antibodies for 2 hours at room temperature.After secondaries, tissue was washed with PBT and stained with DAPI (final concentration 0.5ug/mL) for 15min. The following primary antibodies were used: mouse anti-Hnt (1G9, 1:75), mouse anti-Tgo (1:15), mouse anti-Cut (2B10, 1:15), mouse anti-Br-C (25E9.D7, 1:15) from Developmental Study Hybridoma bank, rabbit anti-GFP (1:4000; Invitrogen), rabbit anti-RFP (1:2000, MBL international), guinea pig anti-Sim (1:1000; a gift from Dr. Stephen Crews, University of North Carolina at Chapel Hill School of Medicine, Chapel Hill, USA). Alexa Fluor488 and Alexa Flour568 goat secondary antibodies (1:1000; Invitrogen) were used.Fluorescent and DIC images were acquired using a Leica TCS SP8 confocal microscope and assembled using Adobe Photoshop software and ImageJ/Fiji software (can be downloaded from https://fiji.sc/).

Mature follicles with egg morphology defects were imaged using Leica DMi8 microscope to quantify Anterior-Posterior and Dorso-Ventral axis length (microns) using ImageJ/Fiji software. The average AP/DV ratio for each genotype was calculated using Excel by dividing the AP-length by the DV-length of each follicle and subsequently calculating the mean ± SD of all individual AP/DV values.

### Statistical Analysis

Statistical tests were carried out using Excel primarily and in some instances Prism 7 (GraphPad, San Diego, CA). Quantification results are displayed as mean ± SD or mean ± SEM as indicated in figure legends. Statistical analysis between two-groups was conducted using two-tailed Student**’**s t-test. For egg distribution assay used to calculate egg laying time, Chi-square analysis was performed to assess significance.

## Acknowledgement

We would like to thank Drs. Stephen Crews, Wu-Min Deng, Allan Spradling, Celeste Berg, Steven Henikoff, Gerald Rubin, Shinya Yamamoto, Takano-Shimizu, Ken Wan, and Liria Masuda-Nakagawa for sharing fly lines and reagents; Bloomington Drosophila Stock Center and Vienna Drosophila Resource Center for fly stocks; and Developmental Studies Hybridoma Bank for antibodies. We thank Drs. Wei Li and Yuping Huang in Dr. Sun’s laboratory for technical support. We appreciate constructive comments from anonymous reviewers. The Leica SP8 confocal microscope is supported by an NIH Award (S10OD016435) to Akiko Nishiyama. JS is supported by the University of Connecticut Start-Up Fund, NIH/National Institute of Child Health and Human Development Grants (R01-HD086175 and R01-HD097206).

